# Haplotype-resolved assembly of the African catfish (*Clarias gariepinus*) provides insights for semi-terrestrial adaptation of airbreathing catfishes

**DOI:** 10.1101/2023.03.23.533919

**Authors:** Julien A. Nguinkal, Yedomon A. B. Zoclanclounon, Ronald M. Brunner, Tom Goldammer

## Abstract

Airbreathing catfishes are a group of stenohaline freshwater fish that can withstand various environmental conditions and farming practices, including the ability to breathe atmospheric oxygen. This unique ability has allowed them to thrive in semi-terrestrial habitats. However, the genomic mechanisms underlying their adaptation to adverse ecological conditions remain poorly understood. Here, we sequenced the genome of the African catfish *Clarias gariepinus*, one of the most commonly farmed clariids. By integrating different long reads sequencing technologies, we generated a chromosome-level assembly with high-resolution haplotypes, including the male-specific haplotype. The diploid assembly yielded 58 contigs spanning 969.72 Mb with a contig N50 of 33.71 Mb. We report 25,655 predicted protein-coding genes and 49.94% repetitive elements in the African catfish genome. Several gene families involved in ion transport, osmoregulation, oxidative stress response, and muscle metabolism were expanded or positively selected in clariids, suggesting a potential role in their transition to terrestrial life. The reported findings expand our understanding of the genomic mechanisms underpinning the resilience and adaptive mechanisms of C. gariepinus to adverse environments. These insights will serve as a valuable resource for future studies in elucidating these unique biological traits in related teleosts and leverage these insights for aquaculture improvement.

## Introduction

The *Clariidae* family, commonly referred to as airbreathing catfish, constitutes a group of freshwater fishes that can thrive out of water for extended periods of time by breathing oxygen from the atmosphere^1, 2^. Some of these facultative air breathers have adapted to terrestrial life by developing the ability to walk on land in sinuous movements using their pectoral fins and a protective mucous layer, helping them to retain moisture^3, 4^. These traits enable them to survive in environments with low oxygen levels or stagnant water, such as mangrove swamps, muddy water or flooded forests, which expand their access to new habitats and food sources^2, 5^. To survive in such environments with changing oxygen tensions, this group of fish has developed a bimodal gas exchange capacity in which the gill extracts oxygen from water, and the accessory respiratory organ extracts it from the air. Their accessory airbreathing organ (ABO) comprises a paired supra-branchial chamber in the gill cavity. This adaptation of clariids to semi-terrestrial environments (amphibious traits) is, however, uncommon among bony fish, as only about 11 distantly related fish genera (out of *∼*2,935)^6^ are considered amphibious with the ability of bimodal respiration^3^. These independent adaptations and traits diversifications are excellent examples of convergent evolution in teleosts. According to FishBase resources, (https://www.fishbase.se/search.php), the *Clariidae* family comprises 16 genera and 116 species, with clariids being the most widespread and diverse group with more than 32 recognized species. Many clariids are well-established aquaculture species, including the African catfish (*Clarias gariepinus*, Burchell, 1822), one of Africa’s most promising endemic aquaculture fish^7^.

*C. garipinus* is found primarily throughout Africa, where it was first introduced in aquaculture around the mid-1970s. This omnivorous fish is quite resilient due to its ability to cope with extreme environmental conditions, tolerate various land-based farming practices and a large diet spectrum^8–11^. In addition to its rapid growth, extreme robustness^11, 12^, and relatively high fecundity, *C. gariepinus* can withstand high levels of ultraviolet B (UV-B) radiation and dramatic temperature fluctuations in non-aquatic environments^13, 14^. This ecological flexibility could explain its hardiness and wide geographical distribution. Interspecies hybridization with closely related clariids has been shown to improve *C. gariepinus* environmental tolerance, manipulate sex ratios, and eventually increase growth performance, making it a highly efficient aquaculture fish^15^. As a result, the African catfish is considered an excellent biological model for studying amphibious traits (i.e., bimodal breathing) and terrestrial transition^16–18^. However, current genomics research has primarily focused on phylogenetic and domestication studies^9, 19–21^, as well as on sex-chromosome and karyotype evolution utilizing only a limited panel of molecular markers^22–24^. *Clarias gariepinus* genome is made up of 2*n* = 2*x* = 56 chromosomes (18 m + 20 sm + 18 st/a)^25^ with a fundamental number (NF) of 94. Its chromosome system has historically been contentious. Previous findings suggested a XX/XY male heterogametic chromosomal system^26–29^, while others pointed to a ZZ/ZW female heterogametic sex determination system (SDS)^30, 31^. However, recent high-throughput sequencing data research has shown that both systems coexist in *C. gariepinus*^22, 23^. The coexistence of both SDSs is most likely heavily influenced by environmental and social factors, as well as geographical habitat: the ZZ/ZW system is indicated in African wild ecotypes^25, 30^, XX/XY system is observed in some anthropogenically introduced populations in Europe and China^27, 28, 32^, and both systems were evidenced within the same population in Thailand^22, 33^.

The lack of genomic resources, including reference genomes, haplotype information, and expression data, has hampered the validation of these SDSs. Yet, few genomic resources of related clariid species, such as the walking catfish (*Clarias batrachus*)^34^ and the Indian catfish (*Clarias magur*)^35^, are publicly available. Despite being only at the scaffold levels and highly fragmented with thousands of gaps, these assemblies provide valuable resources for comparative genomic analyses. However, more high-quality genome data are still needed to advance our understanding of the evolution and adaptation of airbreathing catfish to terrestrial habitats. Gold standard genomes, such as telomere-to-telomere (T2T) and fully phased genomes^36–40^, could facilitate not only studies on sex-chromosome evolution and allele-specific expression but also provide promising tools for investigating biological mechanisms that shape the robustness and adaptation of airbreathing catfishes. T2T genome assembly aims to build a complete and accurate representation of a chromosome from one telomere to the other^41^. T2T assemblies are designed to fill all gaps in a chromosome sequence, including repetitive and complex regions. This includes achieving end-to-end continuity as well as accurately resolving repetitive regions and structural variations. To achieve high-quality T2T assemblies, extensive sequencing technologies, incluing long reads, and a variety of computational methods are frequently used^42, 43^. The identification of alleles that are co-located on the same chromosome is known as haplotype phasing. A fully pahsed genome assembly is one in which the two haplotypes (maternal and paternal) have been separated and assigned to their respective chromosomal sequences^44^. This means that each genomic region is linked to a specific haplotype, allowing for the precise determination of allelic variants and comprehension of haplotype-specific information. Fully phased assemblies are especially useful for examining genetic variations, population genetics, and the inheritance of specific traits^45^.

To better understand the adaptive strategies of air-breathing fish, we carried out genome sequencing and assembly of the African catfish using HiFi PacBio and Nanopore Technologies, as well as long-range phasing information from Hi-C. We derived a near-T2T, fully phased genome assembly of the African catfish. Through a combination of comparative genomic approaches, several genes, biological pathways and processes that are likely associated with the resilience and the emergence of amphibious traits of *Clariidae*, were identified. Our results provide a new genomic basis for functional validation of the molecular mechanisms underlying clariid resilience and their transition out of the water, with potential commercial and ecological implications.

## Methods

### Sample Collection and DNA extraction

Tissue samples, including muscle, liver, and gonads, were collected from one adult male (approximately one-year-old) African catfish in the Experimental Aquaculture Facility of the Research Institute for Farm Animal Biology (Dummertorf, Germany). Prior to tissue collection, the fish was euthanized by immersing it in an overdose of 2-phenoxyethanol (50 mg/L) for 15 minutes, followed by a bleed cut in the head and posterior spinal cord. Tissue samples were immediately frozen in liquid nitrogen and stored at −80◦ C. Following the manufacturers’ standard protocols, we performed genomic DNA extraction using the DNeasy Blood & Tissue Kit (Qiagen) and libraries preparation strategies specific to the sequencing technologies used in this study.

### Libraries preparation and genome sequencing

Genomic DNA (gDNA) sequencing data were generated by different platforms, including Oxford Nanopore (ONT) long reads, PacBio high-fidelity (HiFi) reads, Illumina paired-end reads, and paired-end Hi-C reads (**Figure 1a**). Illumina short-insert (450 bp) libraries were prepared from liver tissues using an Illumina TruSeq Nano DNA Library Prep Kit and paired-end (PE150) sequenced on the Illumina Novaseq 6000 sequencing platform (Illumina, Inc., San Diego, CA, USA). We used gonad tissues for ONT PromethION library preparation and sequencing, following the manufacturer’s (Oxford Nanopore Technologies, Oxford, UK) guidelines. In addition, we sequenced a single flow cell on the PromethION instrument, yielding 84 Gb of data and a sequencing depth of around 80×, with a maximum read length of 330 kb and an N50 of 32 kb. Liver and muscle tissues were pooled for HiFi library preparation and sequenced on the PacBio Sequel IIe sequencing platform (Pacific Biosciences of California, Inc.). We sequenced four SMRT cells, yielding around eight million CCS reads (141 Gb of data) with an N50 of 16 kb and average base call accuracy greater than 99.7%. A Hi-C library was generated using the Arima-HiC kit and following its standard workflow (Arima Genomics, San Diego, CA, USA). All sampled tissues were pooled and then sequenced paired-end (PE150) on an Illumina HiSeq X platform, yielding 182 million read pairs, corresponding to approximately 55× genome coverage. An overview of generated whole-genome sequencing data is provided in the **Supplement Table 1**.

**Figure 1.**
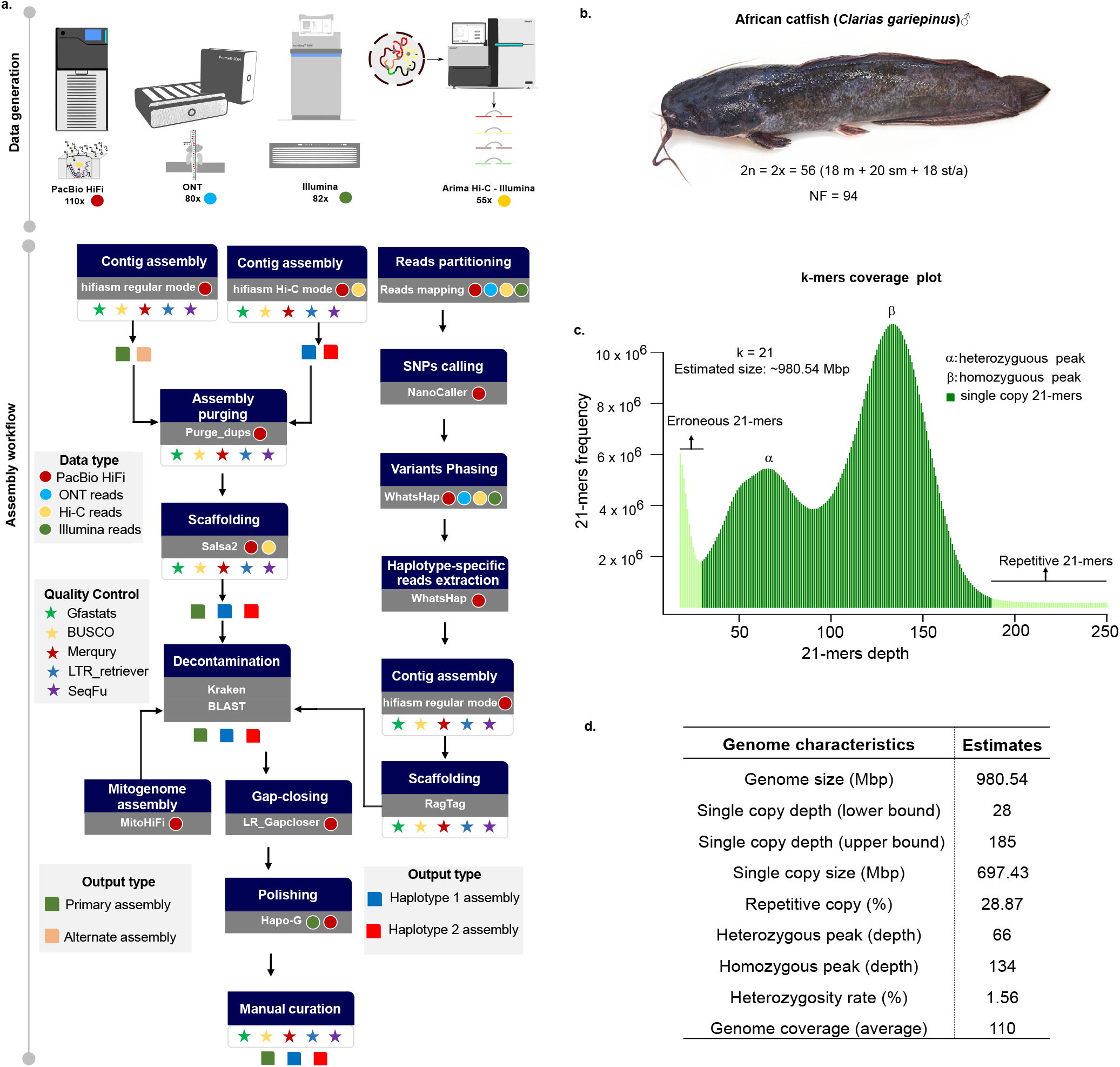
Haplotype-resolved genome assembly workflow of *Clarias gariepinus* and genome survey analysis. **a** The workflow developed to build a haplotype-resolved genome assembly of the African catfish. Generated genomic sequencing data include Illumina paired-end 150, PacBio’s long high-fidelity (HiFi) reads, Oxford Nanopore (ONT) ultra long reads and Hi-C data. A primary assembly and two haplotype-resolved assemblies were obtained using three assembly modes that combined different data types; **b** The African catfish specimen whose genome was sequenced in this study with the chromosome number for male individuals: A diploid genome with 18 metacentric (m), 20 submetacentric (sm), and 18 subtelomeric/acrocentric (st/a) chromosomes. NF is the fundamental number indicating the total number of chromosome arms; **c** *K*-mer frequency distribution of the diploid genome of the African catfish and its size estimate; **d** Preliminary genome characteristics estimated using *k*-mers analysis.

### Genome survey analysis

To estimate the preliminary properties of the African catfish genome, we performed a genome-wide *k*-mer analysis using the k-mer Analysis Toolkit (KAT) (v2.2.0)^46^. Briefly, we generated *k*-mer frequency count (*k* = 21) from high-quality genomic HiFi reads using KAT hist function. With the resulting 21-mer histogram, we used a custum R script (https://shorturl.at/dfnM5) to estimated the genome size, heterozygosity rate, repeat content, and 21-mers derived from errors and sequencing bias. Low-frequency *k*-mers (*depth <* 19) were filtered out. The heterozygosity rate was estimated using the formula: Heterozygosity rate = (Number of distinct k-mers / Total number of k-mers) / 2. We rendered these genomic properties using ggplot2 in R (**Figure 1**).

### Haplotype-resolved chromosome-scale assemblies

Three strategies were used to generate phased assemblies: hifiasm regular mode, HiFi+Hi-C mode, and haplotype-specific HiFi reads obtained through read partitioning. The output assemblies include a primary assembly (Pim), an alternate assembly (Alt), and two haploid assemblies that include haplotype 1 (Hap1) and haplotype 2 (Hap2). The primary assembly is a more contiguous pseudo-haplotype assembly with long alternating stretches of phased blocks that capture both the homozygous regions and a single copy of the heterozygous alleles. Hap1 and Hap2 are phased assemblies representing the entire diploid genome, consisting of parental haplotypes. We used the haplotype-resolved assembler hifiasm (v.0.16.1)^44^ in regular mode (i.e., without Hi-C data) with default parameters to build a contig-level primary and alternate assembly with clean PacBio HiFi reads. Furthermore, a combination of HiFi and PE Hi-C reads was used in hifiasm to generate a set of two haplotype-resolved, phased contig-level (haplotig) assemblies (i.e., hifiasm Hi-C mode). With purge_dups (v1.2.6)^47^, we identified and removed contigs corresponding to haplotypic duplications, false duplications, sequence overlaps, and repeats. To construct chromosome-level phased assemblies, Hi-C PE data were aligned to the purged contigs using a slightly modified Arima Genomics mapping pipeline^48^. This modified pipeline added alignments quality filtering and skipped the second deduplication step. In particular, Alignments with MAPQ scores less than 10 were removed. SALSA2 (v2.3)^49^ performed chromosome scaffolding in three iterations.

Using the phasing information of the haplotype-specific HiFi reads, we generated a set of two haploid assemblies following the workflow described in Garg et al. (2021)^37^. In brief, all genomic reads generated in this study were aligned to the unpolished primary assembly generated in hifiasm regular mode, using minimap2 (v2.2.24)^44^ and BWA-MEM (v0.7.17)^50^ for long and short reads, respectively. We then used HiFi alignments to call heterozygous SNPs using NanoCaller (v.3.0.0)^51^. WhatsHap (v1.4)^52^ was utilzed to phase heterozygous SNPs leveraging inherent phasing information of HiFi, Hi-C, ONT, and Illumina alignments (**Figure 2**). For each genotype, we extracted haplotype-specific HiFi long reads, which were then assembled independently with hifiasm regular mode (**Figure 1a**). High-quality chromosome-scale phased assemblies, including Hap1 and Hap2 were then built using Ragtag (v2.1.0)^53^. The mitogenome was assembled using the mitoHiFi (v2.2)^54^ workflow. To check for putative contaminations, contigs were searched against all RefSeq microbial genomes using Kraken2^55^. In addition, a megaBLAST search against non-animal chromosome-level assemblies from RefSeq was performed, requiring e-value ≤ 10^−5^ and sequence identity ≥ 98%. We applied LR_Gapcloser^56^ with clean HiFi reads to fill unresolved gaps in the Prim assembly. The Hi-C contact maps were visually inspected after polishing and iterative gap-filling to detect potential assembly errors. A few apparent misplacements and orientations of large contigs were identified and manually corrected. The Hapo-G pipeline^57^ was used with default parameters to polish the Prim assembly using PacBio HiFi reads.

**Figure 2.**
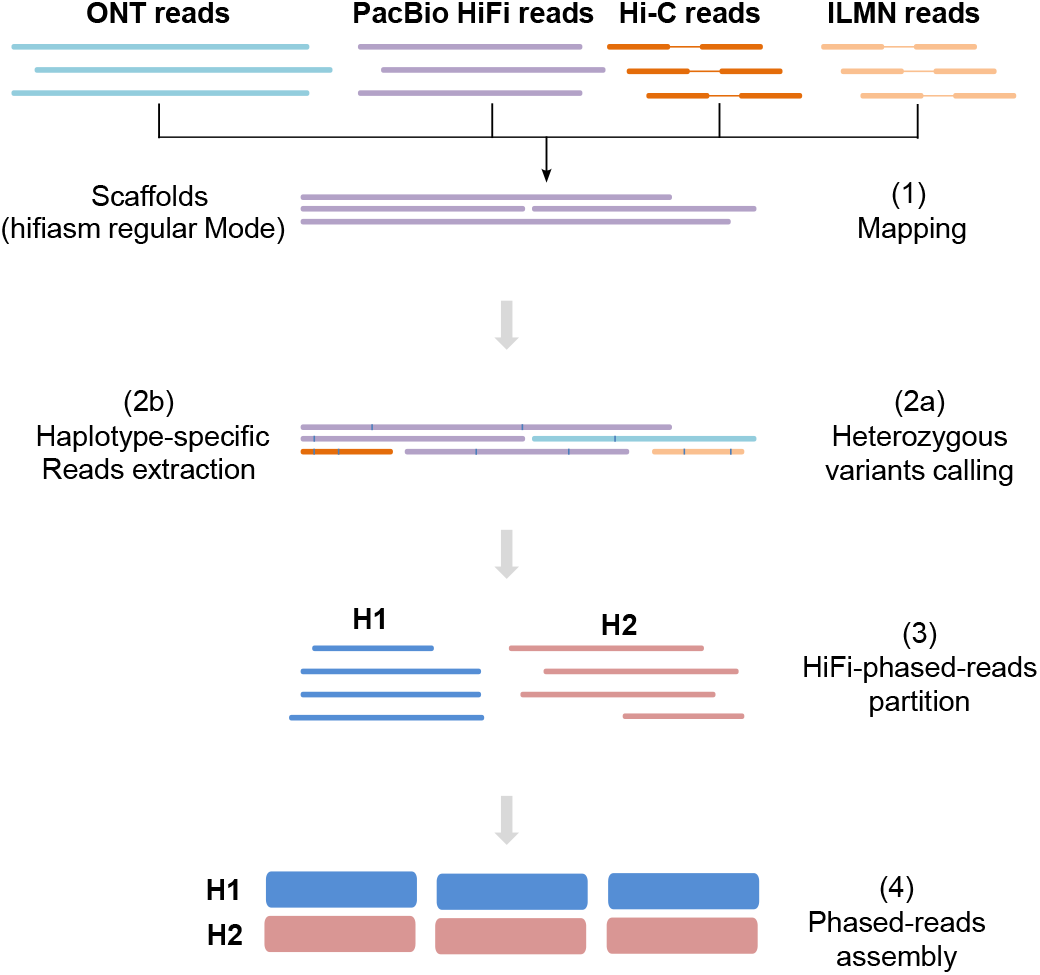
Reads partioning assembly approach. As a reference, the primary assembly obtained in Hifiasm rergular mode was used. After aligning read data from ONT, PacBio HiFi, Hi-C, and Illumina to that reference (1), heterozygous variants were called (2a), and haplotype-specific reads were extracted using WhatsHap (2b). Partitioned reads (3) were then *de novo* assembled into two distinct genome assemblies, one for each haplotype (4).

### Genome assembly quality assessment

Commmon assembly quality metrics were used to assess the overall completeness and accuracy of the African catfish genome assemblies. Benchmarking Universal Single-copy Orthologs (BUSCO) (v5)^58^ with the Actinopterygii dataset and mapping RNA-Seq data from the same species to genome assemblies were conducted to assess gene completeness. The *k*-mer completeness, phasing accuracy, and heterozygosity of the two haplotype assemblies were evaluated by Merqury (v1.3)^59^, using hapmers derived from haplotype-specific reads. The mapping statistics of the raw NGS reads, including Illumina, Hi-C, ONT, and HiFi, were calculated for each assembly. Regarding completeness, phasing accuracy, and contiguity, the haplotype-resolved assembly with HiFi + Hi-C outperformed all other approaches. Unless otherwise stated, we used this assembly in the various subsequent analyses in this study.

### Identification of the putative sex-specific haplotype

To identify the putative paternal haplotype in our assemblies, the full-length nucleotide sequences of two previously identified and validated male-specific DNA markers (Accession numbers: CgaY1: AF332597; CgaY2: AF332598) were obtained from GenBank^29, 60^. Both sequences, 2.6 kb (CgaY1) and 458 bp (CgaY2) in length, were BLASTed against Hap1, Hap2, and Prim assemblies, requiring stringent mapping criteria (identity >98%; query coverage>98%).

### Repeats annotation

Assemblies were annotated independently to avoid a skewed comparison. The methods described here were used to annotate genes and repeats in haplotypes and primary assemblies. RepeatModeler (v2.0.3)^61^ was used to analyze and predict repeat sequences and dependencies such as TRF, RECON, and RepeatScout. Using MITE Tracker^62^, we identified miniature inverted-repeat transposable elements (MITEs). GenomeTools^63^ and LTR_Retriever (v2.9.0)^64^ were used to analyze full-length LTRs. Furthermore, we retrieved all teleost-specific transposable elements (TEs) from FishTEDB^65^, a curated database of TEs identified in complete fish genomes. We used cd-hit (v4.8.1)^66^ to cluster repeat elements with identities greater than 98%. Repeatmasker (v4.1.3)^67^ was used to mask the genome with the final custom non-redundant set of repeats.

### Genes annotation

Protein-coding genes were annotated in *C. gariepinus* genome using *ab initio*, homology-based, and transcriptome-based prediction methods. For homology-based prediction, we obtained high-quality protein sequences from UniProt, which were combined with homologous protein sequences from nine closely related catfish species (**Supplement Table 2**). To map these homologous protein sequences to the African catfish genome, we used TBLASTN with an e-value cutoff of 1e-10. We only kept the highest-scoring alignments with a minimum identity score of more than 80%. The top-scoring proteins were then mapped to the assemblies to predict putative gene models using Exonerate (v2.4.0)^69^. The transcript-based gene prediction was carried out using RNA-Seq data from a conspecific *Clarias gariepinus* individual with available RNA-Seq reads in the Sequence Read Archive (SRA) (BioProject-Accession: PRJNA487132). The quality-filtered reads were mapped to our African catfish assemblies using HISAT2 (v2.2.1)^70^ to detect splice junctions and StringTie2 (v2.2.0)^71^ was then used to assemble transcripts into gene models.

Augustus (v3.4.0)^72^, Genscan^73^, GeneMark-EP^74^, and GlimmerHMM^75^ were used for *ab initio* gene prediction, along with RNA-seq transcript evidence. We used RNA-Seq alignments to train Augustus and GlimmerHMM. In GeneMark-EP and Genscan, we used the default settings. We integrated the genes model prediction from the three methods using the funannotate pipeline (v1.8.13)^76^ to build a consensus, non-redundant gene set. Finally, the resulting gene set was filtered to remove genes with no start or stop codon, an in-frame stop codon, or a coding sequence (CDS) shorter than 180 nucleotides (nt). Genes with high similarity (>90%, e-value < 1e-10) to transposable elements were also removed from the final coding genes set. Several classes of non-coding RNA (ncRNA) genes have also been predicted. tRNAscan-SE^77^ with eukaryote parameters were used to predict transfer RNAs (tRNAs). RNAmmer (v2.1)^78^ was used to identify eukaryotic ribosomal RNA, and the miRDeep2 pipeline^79^ was used to predict putative microRNAs based on homology to eukaryotic mature miRNA sequences in the miRBASE database^80^.

### Functional annotation of protein-coding genes

The functional annotation of protein-coding genes was achieved using BLAST to align predicted protein sequences to RefSeq non-redundant proteins (NR), nucleotides (NT), and UniProtKB/Swiss-Prot databases. Eggnog-mapper (v2.1.9)^81^, and Interproscan (v5.56-89.0)^82^ were used to query BLAST top hits (query_coverage > 60%, identity_score > 80%) to obtain Gene Ontology (GO) annotations and gene names via ortholog transfer.

### Orthologs and phylogenetics analyses

The annotated genome *C. gariepinus* allowed us to understand its evolution and estimate divergence time within catfish species. We downloaded protein sequences from NCBI of 14 catfish species from six lineages, including *Clariidae* (five species), *Ictaluridae* (two species), *Siluridae* (one species), *Pangasiidae* (three species), *Bagridae* (two species), and *Sisoridae* (one species). **Supplement Table 2** contains extensive meta-information on these species. Throughout this analysis, two *Cyprinidae* species were used as outgroups: the goldfish (*Carassius auratus*) and the common carp (*Cyprinus carpio*).

Gene families from the 14 catfish, including outgroup species were identified using the OrthoFinder pipeline with default settings, except that the *diamond_more_sensitive* flag was set in alignment parameters. In brief, an all-vs. all BLASTP comparison with an e-value threshold of 1 × 10^−10^ was performed with all proteins and then genes were clustered into orthogroups using the MCL algorithm. The coding sequences of the single-copy orthogroups were aligned with mafft and concatenated into a super gene for each species. The rooted species tree and gene trees were inferred using single-copy orthologs. The MEGA11^83^ program for Linux was used to estimate the divergence times among the species using rapid relaxed-clock methods^84^ and molecular clock data for calibration constraints obtained from the TimeTree database^85^ between the black bullhead (*Ameiurus melas*) and the goonch (*Bagarius yarrell*).

### Gene families evolutionary analysis

The Computational Analysis of Gene Family Evolution (CAFE) analysis was performed with default parameters to estimate the contraction and expansion of gene families for the 14 catfish species mentioned above. In brief, the time-calibrated ultrametric species tree and orthologous gene families were sent to CAFE (v5)^86^, and significant (*p*−*value* < 0.05) size variance of gene family expansions and contractions were identified using 1000 random samples, and deviated branches were determined using the Viterbi algorithm implemented in CAFE with a branch-specific p-value less than 0.05. A custom bash script was used to identify significant species-specific gene gain or loss in gene families. Finally, we used the KOBAS-i tool^87^ to perform functional enrichment analyses and to identify pathways and GO terms significantly associated with gene families expansion in the airbreathing catfishes examined in this study.

### Positive selection analyses

We used the PosiGene pipeline^88^ to scan genome-wide positive selection among the aforementioned catfishes, detect selective signatures and understand their role in the adaptive mechanisms of amphibious airbreathing catfishes (*Clariidae*). Positive selection in the *Clariidae* branch was scanned using branch-site tests based on one-to-one single-copy orthologs. The yellow catfish (*Tachysurus fulvidraco*) served as an anchor species, while the black bullhead and goonch served as outgroups. The false discovery rate (FDR) threshold for significantly positively selected genes was set to less than 0.05.

### Gene duplication events analysis

We examined ten catfish with chromosomal-level genome assembly to identify different types of gene duplication events that could have shaped their evolution. We identified gene pairs derived from whole-genome (WGD), tandem (TD), proximal (PD), transposed (TRD), or dispersed (DSD) duplications using the workflow described by Qiao et al (2019)^89^ and the DupGen_finder pipeline (https://github.com/qiao-xin/DupGen_finder). For each duplicate gene pair, we calculated the synonymous (Ks) and non-synonymous (Ka) nucleotide substitution rates between the two paralogs using the calculate_Ka_Ks_pipeline^89, 90^.

## Results

### Whole genome sequencing

Sequencing and assembly of teleosts genomes are complicated due to inherent heterozygosity, retained ohnologs, and high repeat content. This study used a stepwise data integration and assembly validation approach with four complementary NGS technologies to generate the African catfish’s haplotype-resolved assembly. We sequenced tissues from a male *C. gariepinus* specimen (**Figure 1b**) using Illumina PE reads (*∼*82×), PacBio’s HiFi reads (*∼*110×), Oxford nanopore reads (*∼*80×), and Hi-C library sequencing data (*∼*55×) (**Figure 1a**). We used 120 Gb of high-quality HiFi data to conduct genome survey analysis. The *k*-mer analysis (*k* = 21) revealed an estimated genome size of *∼*980 Mbp, a relatively high heterozygosity rate of 2.12%, and the expected repetitive sequences accounted for approximately 46% of the entire genome (**Figure 1c-d**). The *k*-mer spectra histogram illustrates the high heterozygosity between both haplotypes, with homozygous regions consisting mainly of 2-copy *k*-mers and heterozygous regions consisting mostly of 1-copy *k*-mers, as expected from a diploid genome (**Figure 1c**).

The *C. gariepinus* genome was *de novo* assembled using three methods: the standard HiFi-only mode, HiFi+Hi-C mode, and reads partitioning using SNPs phasing information from HiFi, Illumina PE, Hi-C and ONT sequencing data. Except for the HiFi-only mode, which produced a partially phased assembly consisting of a collapsed primary assembly and an incomplete and fragmented alternate assembly, we benchmarked the contiguity and phasing accuracy of haplotypes from both HiFi+Hi-C and reads partitioning approaches. Although there was only a slight difference in assembly contiguity and structural accuracy between the two methods, the assembly obtained with HiFi+Hi-C reached a slightly better accuracy (**Supplement Table 3**). Here, we present the HiFi+Hi-C assembly, which has been extensively validated and used as a reference assembly for the various analyses performed throughout this study.

### Phased assemblies of the African catfish genome

Following QC filtering and duplicates removal, the initial phased contig-level assembly yielded 58, 142, and 212 sequences, with contigs N50 values of 33.71 Mb, 32.12 Mb, and 19.53 Mb for the Primary, Haplotype-1, and Haplotype-2, respectively. As confirmed later by scaffolding with Hi-C data, more than half (*n* = 34) of the 58 primary contigs represented already entire chromosomes end-to-end or full-length chromosome arms. After polishing and quality improvement, enhanced fully phased chromosome-scale assemblies were obtained by scaffolding contigs into 28 chromosomes and filling most gaps. The Primary assembly (Prim) chromosomes were sorted and numbered in order of decreasing physical size. Synteny mapping to Prim was used to number Haplotype-1 (Hap1) and Haplotype-2 (Hap2) chromosomes. The chromosome sizes range from 52 Mbp (chr1) to 21 Mbp (chr28), with a median length of 32.3 Mbp. The high heterozygosity rate (1.56%) of the African catfish genome may have facilitated this successful haplotype separation, as it has previously been shown that higher heterozygosity rate aids efficient genome unzipping^59^. The evolution of assmebly metric after each processing stage is depicted in **Supplemnet file S1**.

Approximately 99% of the assembled genome is spanned by the 28 chromosomes of the Primary assembly, which have no gaps, whereas Hap1 and Hap2 contained only 0.01% and 1.44% unresolved nucleotides (gaps), respectively, mostly in repeat-rich genomic regions. Hi-C analysis identified four chimeric contigs, which were manually examined and corrected. The final haplotype-resolved assembly size for Prim, Hap1 and Hap2 is 969.72 Mb, 972.60 Mb, and 954.24 Mb, respectively. Only Hap2 dramatically increased the N50 metric from 19 Mb to more than 33 Mb at the scaffold level (**Table 1**). Chromosome-wide analysis of telomeric repeats captured the terminal and tandemly repeated motif (*TTAGGG/CCCTAA*)*_n_* at both chromosomal termini (first and last 25 kbp window) in 21 of 28 *C. gariepinus* chromosomes (**Figure 3a**). Terminal telomeric repeats captured in the first and last 25 kbp windows range from 300 bp to 14 kbp, with an average length of 4.5 kbp (**Supplement Table 4**). Extending the search window to 1 Mbp did not result in a significantly larger copy number of terminal telomeric repeat. Terminal 25-kbp windows had significantly (p.adjust < 0.01) larger telomere sizes and densities per kbp than terminal 1 Mbp windows (**Figure 3b-c**). This result suggests that the terminal 25 kbp windows captured the majority of full-length telomeric repeats in our African catfish chromosomes assembly, which is consistent with previous findings indicating that the length of telomeric DNA in fish ranges from 2 to 25 kb^91–93^. We also identified a few internal telomeric sequences with high copy numbers (*n >* 200). These interstitial or pericentromeric telomeric sequences (ITS) have been evidenced as relics of genome rearrangements in some vertebrates species.

**Figure 3.**
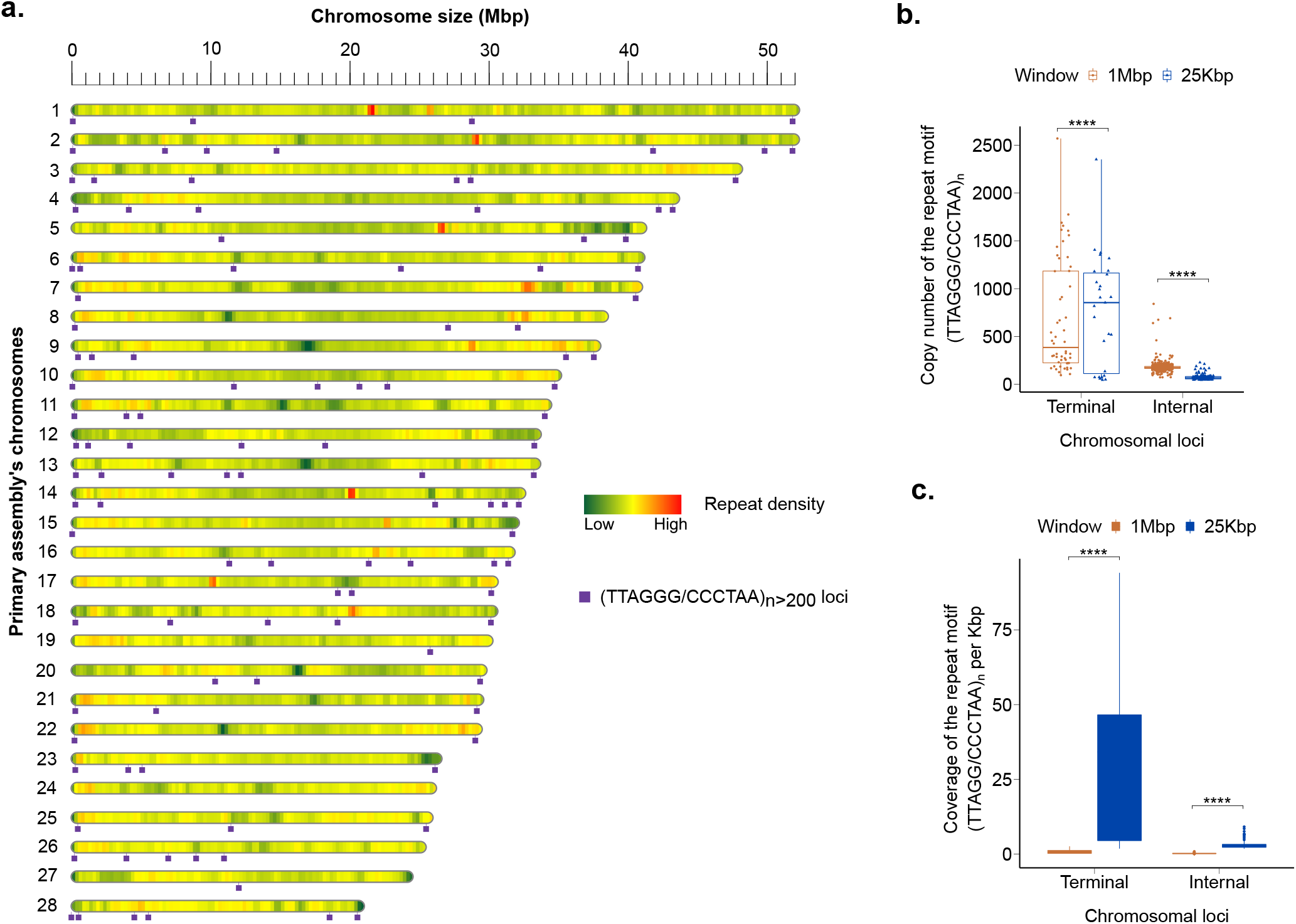
Genome-wide telomere portrait of *Clarias gariepinus*. **a** The purple boxes are chromosomal loci of the tandemly repeated telomeric motif (*TTAGGG/CCCTAA*)*_n>_*_200_ in the Primary assembly. Only telomeric repeats with a minimum size of 1200 bp are shown. The heatmap shows the chromosome-wide repeat density in non-overlapping 500 kbp windows; **b** Boxplots show the copy number distribution of the telomeric repeat motifs (*TTAGGG/CCCTAA*)*_n>_*_45_ in Terminal and Internal 25 kbp and 1 Mbp windows. **** are statistical significance levels of the T-test (*p*−*value <* 0.0001)); **c** Boxplots depict the density of (*TTAGGG/CCCTAA*)*_n>_*_45_ motif per 1000 bp in Terminal and Internal 25 kbp and 1 Mbp windows.

**Table 1.**
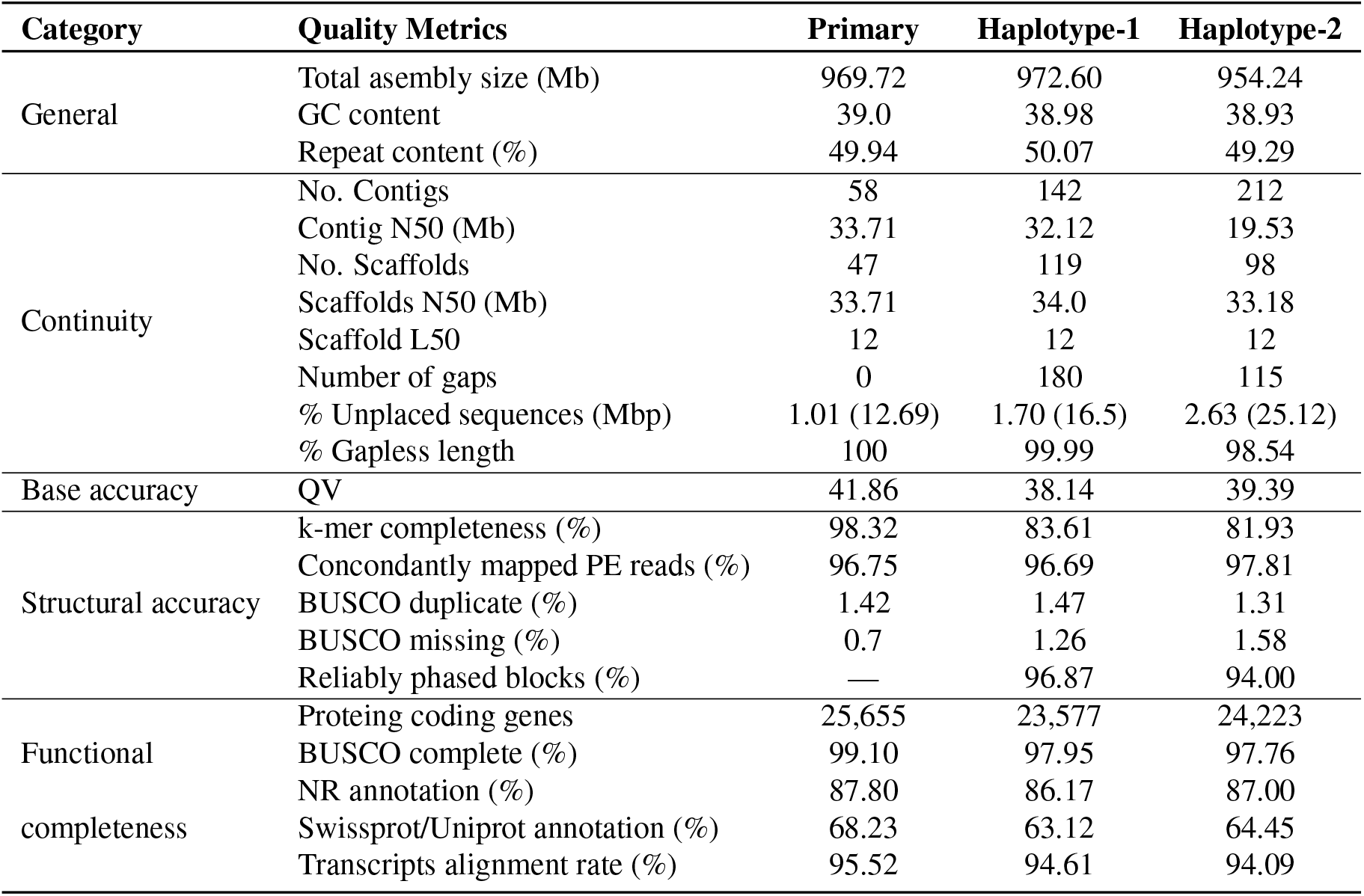
Summary of assembly metric of the *Clarias gariepinus* genome, including the primary (Prim), haplotype-1 (Hap1) and haplotype-2 (Hap2).

### Validation of the male-specific marker CgaY1 (AF332598) on Hap1

CgaY1 (AF332598)^29^, a previously identified male-specific marker in C. gariepinus, was mapped to only one chromosome in Haplotype-1 and Primary assemblies (identity > 99.14; query Coverage > 96.5; e-value = 0). We found no significant hits on Haplotype-2. CgaY1 is on chromosome 24 at position chr24:20208252-20208717 (Prim) and chr24:20319406-20319871 (Hap1). To confirm the absence of this male-specific marker on Hap2, we extracted its flaking sequence (2kb upstream and downstream) and aligned it to chromosome 24 in both the Hap2 and Prim assemblies. We found a single (> 95%) match on Prim but none on Hap2. Although we cannot conclusively determine the Y/W chromosome from this data, we assume that haplotype-1 assembly corresponds most likely to the male-specific haplotype. The Genbank accession numbers for the Primary, Haplotype-1, and Haplotype-2 assemblies are GCA_024256425.1, GCA_024256435.1, and GCA_024256465.1, respectively.

### Genome structural and functional annotation

Integrating ab initio predictions, proteins, and RNA-Seq alignments, we independently annotated the primary assembly and both haploid assemblies. In the collapsed diploid assembly, a total of 25,655 protein-coding gene models were predicted. Hap1 and Hap2 yielded a slightly lower number of predicted genes, with 23,577 and 24,223, respectively (**Table 1**). Approximately 200 genes predicted in Prim were completely missing from Hap1 and Hap2 assemblies. The primary assembly consistently resulted in a better functional annotation, which is to be expected given that the diploid assembly includes both haplotypes and maps a complete representation of the genome structure. Overall, 87.80% of the 73,455 high-quality proteins across the primary assembly and both haplotypes were assigned a function in at least one of the functional databases searched through sequence homology or orthologs mapping. (**Table 1**).

Repetitive sequences made up half (49.94%) of the *C. gariepinus* genome, which roughly corresponded to the estimated repeat content based on *k*-mers analysis. This relatively high repeat content in the African catfish genome is comparable to that found in other catfishes, including *Clarias magur* (43.72%)^35^, *Clarias macrocephalus* (38.28%)^94^, *Pangasianodon hypophthalmus* (42.10%)^95^ and *Hemibagrus wyckioides* (40.12%)^96^. Still, this is still higher than in *Clarias batrachus* (30.30%)^34^, which has a smaller genome size (821.85 Mb). Interspersed repeats are the most abundant class of repetitive elements (46%). Retroelements and DNA transposons accounted for only 12 and 6 per cent of the repeatome, respectively (**Supplement Table 5**). The distribution of genes and repeats across the chromosomes followed the typical distribution in vertebrate genomes, with higher gene densities in GC-rich regions and lower gene densities in repeat-rich distal and pericentromeric regions (**Figure 4**).

**Figure 4.**
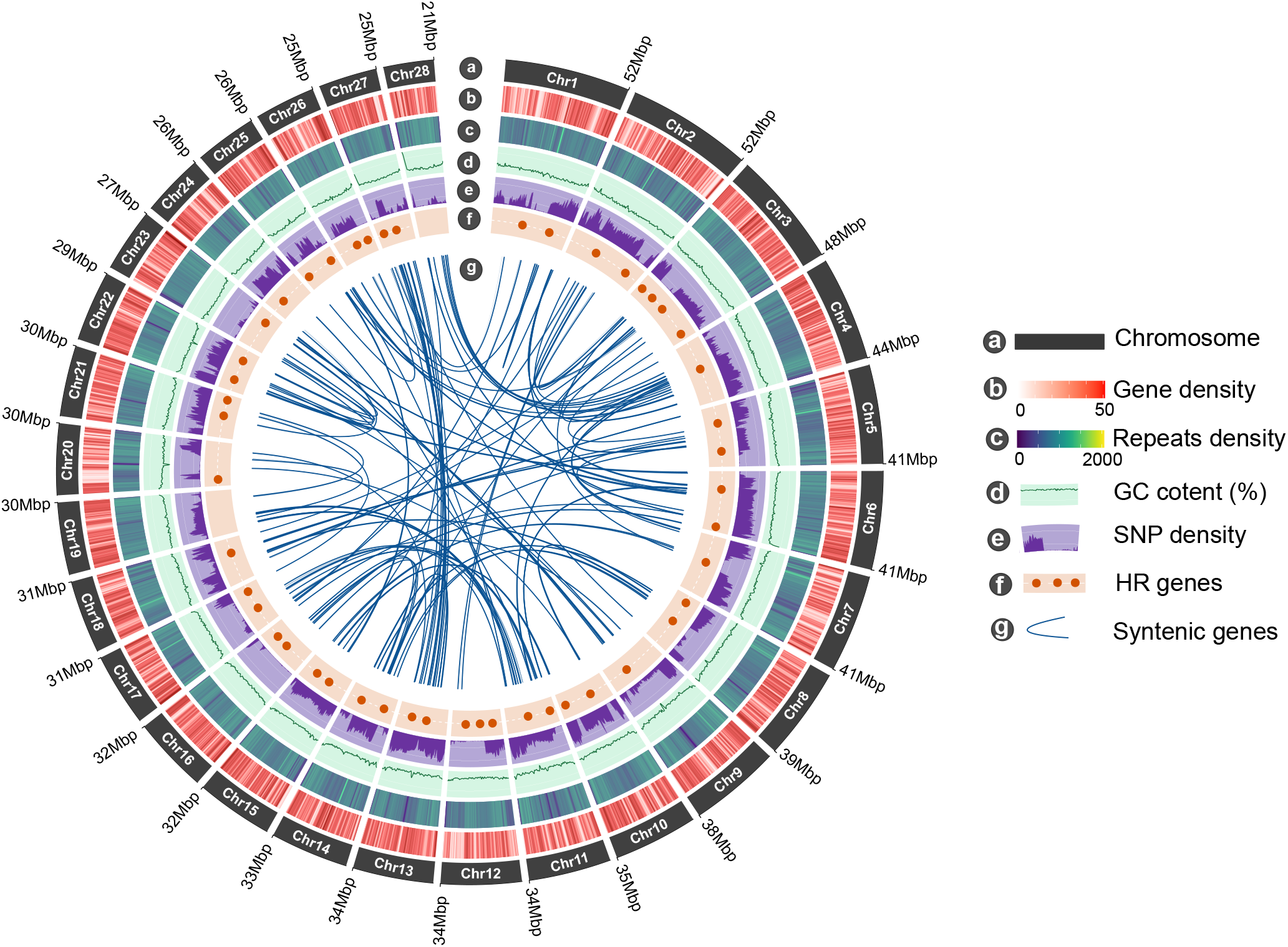
Genomic features of *Clarias gariepinus*. From the outer to the inner circle: **a** Length of the 28 diploid chromosomes (Mb); **b** Chromosome-wide gene density per non-overlapping 500 kb windows; **c** Repeats density in non-overlapping 500 kb windows; **d** GC content; **e** Distribution of heterozygous SNPs density; **f** Chromosomal loci of hypoxia-responsive (HR) genes predicted in the *C. gariepinus* genome; **g** The inner curve lines indicate syntenic gene pairs identified between *C. gariepinus* chromosomes.

Furthermore, we annotated 6,403 full-length ribosomal RNA, 154 microRNA, and 13,536 transfer RNA throughout the African catfish genome. Remarkably, 96% (6150/6406) of the predicted 5S rRNA genes were all found in a single cluster on a 2-Mbp region on both chromosome 4 (*n* = 2455) and chromosome 13 (*n* = 3725). Similarly, 84% (21/25) of the predicted 18S rRNA genes were clustered within the first 500 kbp upstream in the terminal telomeric region of chromosome 27 (**Supplement Figure S1**). This result is consistent with earlier findings^25^ in which 5S rDNA was hybridized on a single site on two *C. gariepinus* chromosomes and 18S on just one site on a submetacentric (sm) chromosome (**Supplement Figure S1**). The ribosomal 18S DNA probe did, in fact, hybridize with the sub-telomeric/telomeric region of a medium-sized sm chromosomal pair in *C. gariepinus*, which most likely corresponds to the 500 kb telomeric region on chromosome 27 in this study. The 5S rDNA sequences were identified as a single hotspot in two subtelomeric/acrocentric (st/a) chromosome pairs in *C. gariepinus*, which is most likely the 2 Mbp large 5S rRNA genes cluster our study evidenced on chromosome 4 and chromosome 13 (in the regions 16-18 Mbp) (**Supplement Figure S1**).

### Assembly assessment and validation

We performed various assessments to validate the high-quality and completeness of our haplotype-phased African catfish genome assembly, including gene completeness, full-length transcript coverage, read mappability rate, phasing accuracy, and genomic k-mer completeness. The BUSCO completeness (99.10%) was comparable between haplotypes and the primary assemblies. Since we missed only 0.7% of the expected universal orthologs, we assume that the gene space spanned by our genome assembly is nearly complete (**Table 1**). Furthermore, approximately 92% of the *C. gariepinus* transcripts could map on our assemblies (> 90% coverage and >90% identity), indicating their high functional completeness. We also mapped genomic reads to our assemblies to assess structural accuracy and found that more than 96.69% of raw PE reads were concordantly aligned. The alignment rate of ONT, HiFi, and Hi-C reads to the primary assembly was 99.91%, 99.95%, and 100%, respectively. The mapping rates to Hap1 and Hap2 were greater than 99% (**Supplement Table 6**). Utilizing the telomere identification toolkit (tidk)^68^, we scanned *C. gariepinus* genome for terminal telomeric repeats (5’-TTAGGG-3’)*_n_* with a minimum length of 270 bp (*n* = 45) in 25 kb windows of chromosomal termini. To be termed ’terminal telomeric repeats’, we required the motif (*TTAGGG/CCCTAA*)*_n_* to exhibit the highest density per 25 kb in the terminal 25 kb windows compared to internal 25 kb windows. All non-terminal telomeric repeats are referred to as internal or interstitial telomeric sequences (ITS).

Merqury assessed the assembly qualities by evaluating phasing consistency and accuracy with haplotype-specific *k*-mers. We expected the sets of haplotype-specific *k*-mers to be entirely distinct for a perfectly phased assembly, with no mixture of k-mers from both haplotypes. Our data shows that Hap1 and Hap2 are orthogonal with only very few haplotype switches and nearly no contamination (**Figure 5a**). Interestingly, homozygous *k*-mers between both haplotypes were ideally shared in the 2-copy peak. In contrast, a substantial amount of haplotype-specific (heterozygous) *k*-mers was distinct in the 1-copy peak in the spectrum copy number plot (**Figure 5b**). The imbalance of *k*-mers specific to each haplotype representing heterozygous alleles is most likely due to significant differences in sex chromosome sizes.

**Figure 5.**
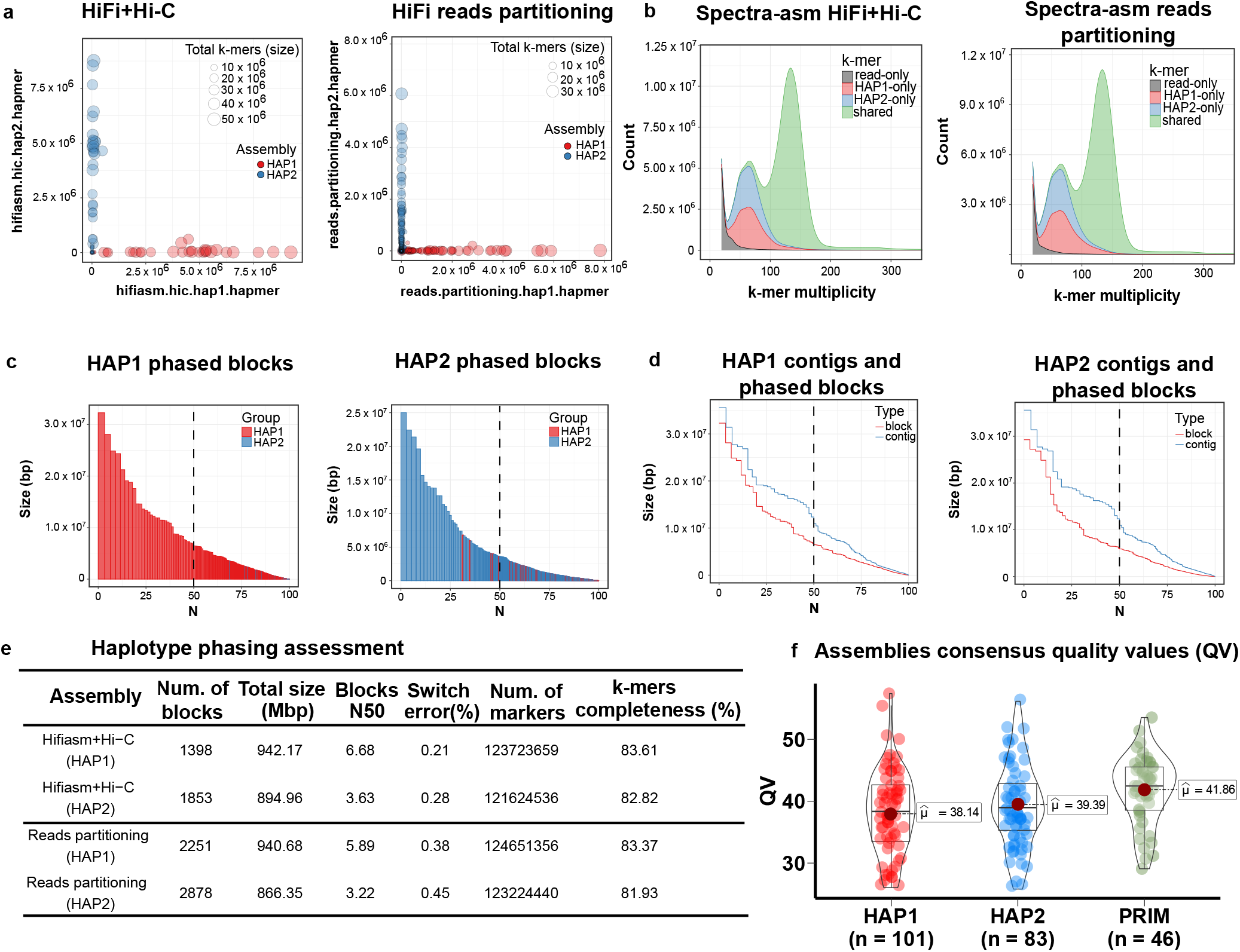
QC plots for evaluating haplotype phasing accuracy, genome contiguity and completeness. **a** Hap-mers blob plot of the Hifi+Hi-C (left) and HiFi reads partitioning assembly (rigth). Red blobs represent HAP1-specific *k*-mers, while blue blobs are the HAP2-specific *k*-mers. Blob size is proportional to chromosome size. A well-phased assembly should have orthogonal hapmers (e.g. HAP1 and HAP2 lie along axis, respectively). Both assemblies show nearly no haplotypes mixture;**b** Spectra-asm plot of HiFi+Hi-C (left) and Reads partitioning (rigth) assemblies. The 1-copy k-mers representing the heterozygous alleles are specific to each haplotype assembly (HAP1 and HAP2), and the 2-copy k-mers, which are only found in the diploid genome, are shared by both assemblies (green). There is no discernible difference between the two assembly approaches. Low-copy *k*-mers (depth < 18) arising from contamination or sequencing errors were removed from the visualization; **c** Phased blocks N* plots of HAP1 (left) and HAP2 (right) assembly, sorted by size. X-axis represents the percentage of the assembly size (*) covered by phased blocks of this size or larger (Y-axis). Blocks from the incorrect haplotype (haplotype switches) are very small and almost entirely absent in the other haplotype. In both haploytpes, more than 75% of the assembly is spanned by phased blocks larger than 1 Mbp; **d** Phase block and contig N* plots showing the relative continuity of HAP1 (left) and HAP2 (right); **e** Statistics for haplotype phasing with switch errors and phased blocks allowing up to 100 switches within 20 kbp; **f** The average consensus quality (QV) distribution for each assembly. Each dot represents a scaffold in the associated assembly.

In our haplotype-resolved genome assembly, the phased blocks originating from the wrong haplotype were tiny and almost entirely absent when plotting them sorted by size (**Figure 5c**). Moreover, the total phased block sizes accounted for 97% and 94% of Hap1 and Hap2 assemblies, respectively. Merqury reported N50 phase block sizes of 3.6 Mbp and 5.5 Mbp with only 0.28% switch error rate when a maximum of 100 consecutive switches were allowed within a 20 kbp window ((**Figure 5d-e**). The collapsed diploid assembly (Prim) recovered 98.32% of the *k*-mers derived from genomic reads, while the haploid assemblies (Hap1 and Hap2) recovered 83.67% and 82.82%, respectively, demonstrating higher genome completeness **Figure 5e**). The average base-level accuracy in the Prim assembly was roughly QV42, corresponding to a rate of less than 0.01%. Hap1 and Hap2 had a slightly lower accuracy than QV40. It should be noted that haplotype assemblies were not polished to avoid introducing more switch errors and a biased homozygosity (**Figure 5f**).

Overall, our assembly quality metrics indicate a gapless, fully phased, and near-T2T assembly of the African catfish genome. Most of these metrics meet or exceed the minimum quality standards^48^ of the VGP consortium. Our reported genome, for example, meets the *c.c.P5.Q42.C98* VGP standard, with *c.c.Pc.Q60.C100*^48^ being the highest standard for finished and gapless T2T vertebrate genomes, such as the recently completed gapless human genome sequence^41^. To the best of our knowledge, this assembly is the first T2T, haplotype-resolved, and the most complete *Siluriformes* (catfish) genome assembly published to date (**Supplement Table 2**).

### Phylogeny, divergence time and evolutionary history of catfishes

The comparative phylogenomic analyses performed with OrthoFinder assigned 336,681 (94%) of 390,198 genes to 27,587 orthogroups shared among catfishes and two outgroup species (common carp and goldfish). A total of 16,281 genes in *C. gariepinus* were found to be orthologous between the 14 catfish species, with 378 being single-copy orthologs. The alignments of single-copy orthologs were used to infer the species tree and evolutionary divergence time (**Figure 6**). Eighty orthogroups comprised 840 genes unique to all airbreathing catfish species, with 208 genes specific to *C. gariepinus* and spanning 80 orthogroups (**Supplement Table 7)**. The vast majority of *C. gariepinus*-specific genes were not characterized in functional databases. Though ten genes belong to the actin family, eight to the peptidase C13 family, and five to the zinc-finger protein family. According to our estimated phylogenetic tree using protein sequences of all homologous single-copy genes, airbreathing catfishes (*Clariidae* clade) split as a monophyletic group around 98 Mya, which is roughly comparable to the divergence time between rodents and humans (96 Mya) (**Figure 6**).

**Figure 6.**
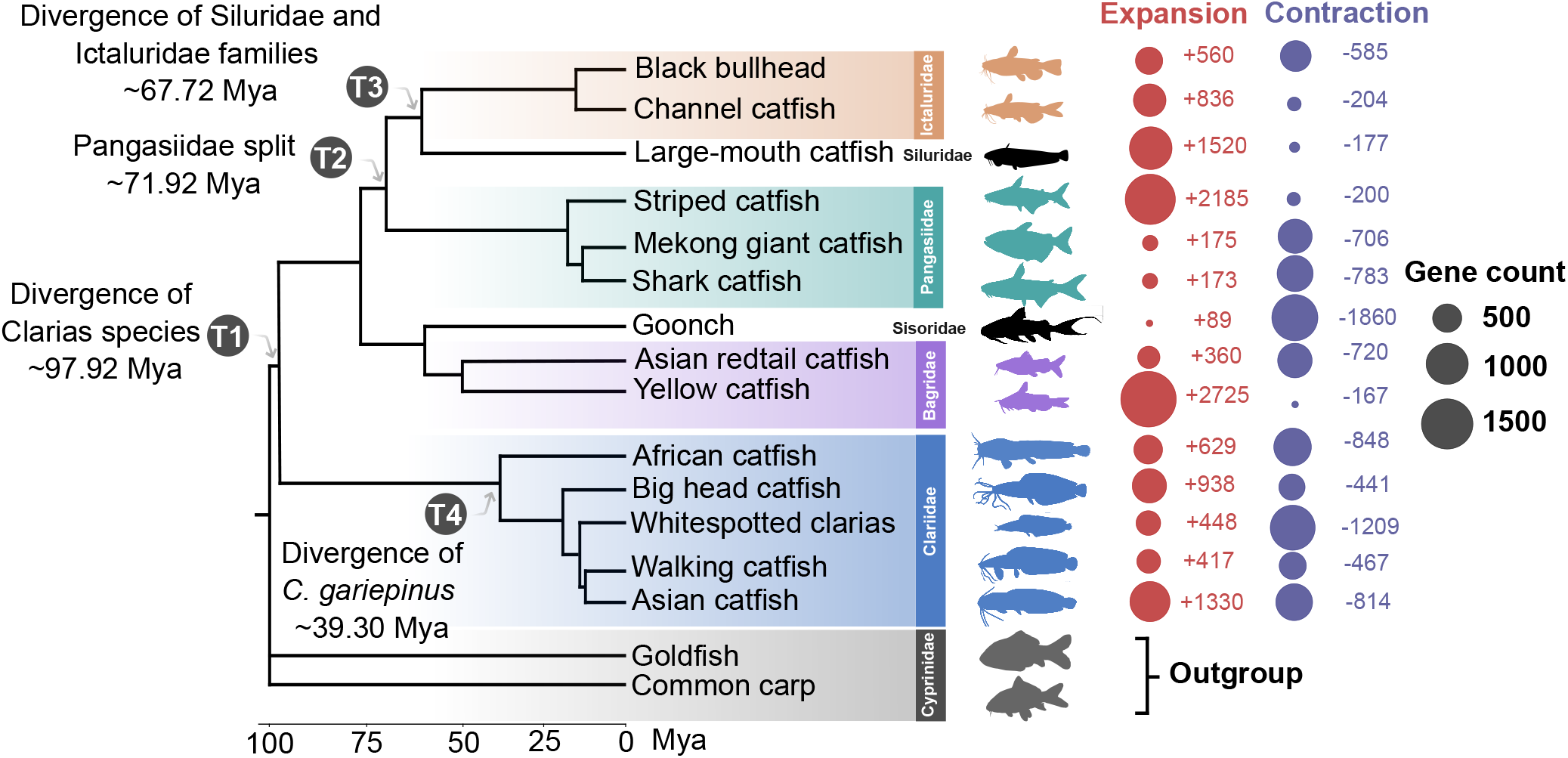
Phylogenomic relationships of major groups of catfishes (Silurinormes). Time-calibrated phylogenetic tree of 14 catfish species based on 1:1 single-copy orthologous proteins. Estimated divergence time as well as the time scale in million years (mya) are shown at the bottom axis. The bubble chart at the right end of the species represents the proportion of gene families that underwent expansion (red) or contraction (blue) in a specific branch. The circle radius is proportional to the number of genes assigned to each category.

The African catfish diverged from clariids last common ancestor (LCA) about 39.3 million years ago, which is consistent with the current understanding of the historical and geographical distribution of the Clariids, with *Clarias gariepinus* being the only clariid species (in our study) native to Africa^97^. In contrast, the other Clarias species are all endemic to Asia. This result aligns with the paraphyly hypothesis previously put up for the genus Clarias^98^. Due to biogeography and adaptive responses to environmental stressors, the African catfish gradually acquired unique traits and features following the split between the African and Asian *Clarias*^99^. Our phylogeny analysis suggests that the Asian *Clarias* clade underwent its first speciation event about 25 Mya, which is consistent with the age of the fossil records available for these species^100^.

### Comparative gene family evolution of airbreathing catfishes

The expansion and contraction of gene families can play an essential role in the adaptation of catfish and other organisms to specific environments by enabling the development and expression of beneficial traits while decreasing the expression of less essential ones. Gene expansion and contraction can lead to phenotypes’ potential gain or loss. To investigate the lineage-specific adaptation of *Clarias*, we used CAFE (Computational Analysis of Gene Family Evolution) to estimate gene family expansions and contractions among 27,587 ortholog groups shared by catfishes, including five airbreathing and nine non-airbreathing catfishes (**Methods**).

We found 1,429 and 2,547 gene families that are significantly expanded or contracted in airbreathing catfish. Gene families in *Clarias magur* had the most gene expansion events (1,330), while gene families in *Clarias fuscus* exhibited the most contraction (1,209) events (**Figure 6**). We identified 629 expanded and 848 contracted gene families in the *Clarias gariepinus* genome. The egalitarian nine homolog gene family (EGLN), the rhodopsin (RHO) gene family, the ferretin (FTH) gene family, and the Carboxypeptidase A (CPA) gene family are some examples of expanded gene families in *C. garipeinus* and other *Clarias* sp. (**Figure 7**). These gene families were all thought to be involved in the environmental adaptation of *Clarias magur*, a closely related species to *C. gariepinus*^35^. The EGLN gene family encodes for prolyl hydroxylase enzymes, which regulate hypoxia-inducible factor (HIF). HIF is a protein that plays a key role in the body’s response to low oxygen levels, and prolyl hydroxylase enzymes regulate HIF expression. The duplication of the RHO gene has been proposed as a mechanism for the adaptation of tetrapods^101^ and amphibious fishes^102–104^ to terrestrial environments. The expansion of this gene family in Clarias may suggest a critical role in their visual system and light adaptation out of water. Finally, FTH proteins have been associated with iron metabolism and are involved in environment-fish-cross-talk^105, 106^.

**Figure 7.**
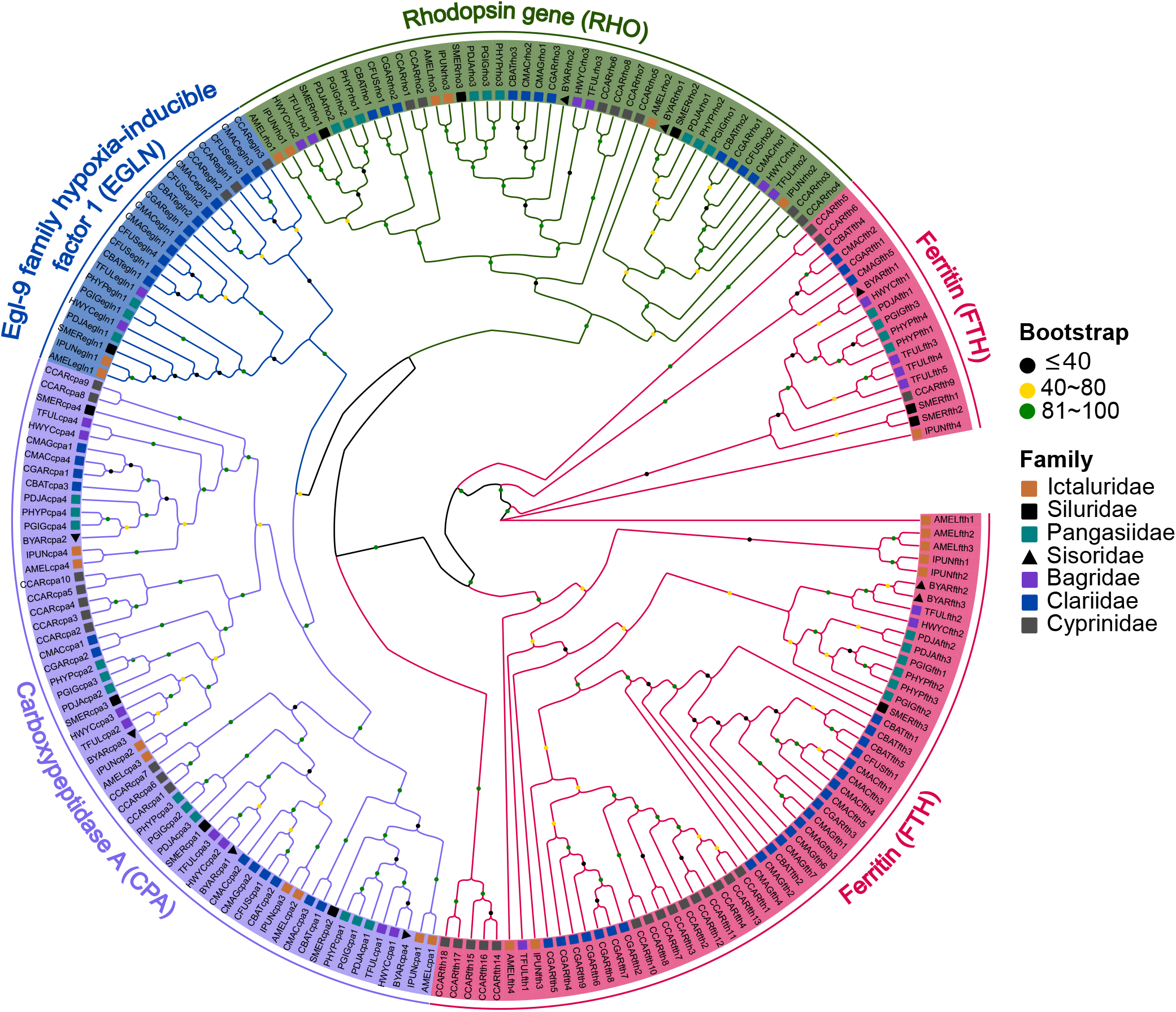
Examples of Clarias-specific expanded gene families. Maximum likelihood gene tree showing the phylogenetic relationship of four gene families significantly expanded only in *Clariidae* (airbreathing catfishes), but not in non-airbreathing catfishes. Species of the same taxonomic family have the same shape and color. Bootstrap values are indicated with black, yellow and green colors.

Expanded gene families in *Clarias* are primarily enriched with ion metal binding, apelin signalling, adrenergic signalling in cardiomyocytes, and neuroactive ligand-receptor interaction pathways, to name only a few. Nucleotide-binding (GO:0000166), anatomical structure development (GO:0048856), response to stimulus (GO:0050896), and cytoskeletal motor activity (GO:0003774) are some of the significantly overrepresented GO terms associated with expanded gene families in these facultative airbreathing freshwater fishes (**Supplement Figures S2-S4**). Overall, gene family expansion in airbreathing catfishes is primarily characterized by expanding gene families encoding for ion transporters and enzymes involved in osmoregulation, metabolism, and energy production. The expansion of these gene families may help airbreathing catfishes cope with the challenges of terrestrial life, such as fluctuating oxygen levels and adapting to new energy sources. The expansion of many gene families involved in cytoskeletal motor activity and anatomical structure development may cause adaptive changes in genes expression to promote the development or modification of specialized anatomical structures, such as gills, labyrinth, blood vessels, and muscles, as well as traits required for low-oxygen environments and efficient terrestrial locomotion and survival.

### Positive selection in airbreathing Clarias

The genome-wide screening for positive selection in airbreathing catfish detected nine protein-coding genes under selective pressure (*FDR <* 0.05) when compared to non-airbreathing catfishes (**Supplement Table 8**). For example, the 3-hydroxybutyrate dehydrogenase (BDH1), a member of the short-chain dehydrogenases/reductases (SDR) protein family found in airbreathing catfishes, accumulated up to 14 conserved non-synonymous amino acid substitutions (sites) across *Clarias* species but not in non-airbreathing catfishes. SDR enzymes are known to be involved in the metabolism of lipids and regulating energy balance^107^, which could be important for airbreathing catfishes to preserve energy balance when moving on land. Additionally, some SDR enzymes detoxify harmful compounds such as pollutants and oxidants in terrestrial environments^108^, which can help airbreathing catfishes survive in these harsh conditions.

### Landscape of gene duplications in catfishes

Gene duplication is most likely another driver of airbreathing catfish adaptation. This process can result in the evolution of new genes and the expansion of gene families, which contribute to the acquisition of evolutionary novelty. Among the 25,655 coding genes in the African catfish genome, 13,809 were derived from diverse gene duplication events. Based on their duplication mode, DupGen_finder (**Methods**) classified duplicated genes into five categories: (i) 496 whole-genome duplicates (WGDs, 3.6%), (ii) 1,463 tandem duplicates (TDs, 10.6%), (iii) 572 proximal duplicates (PDs, 4.14%), (iv) 2,970 transposed duplicates (TRDs, 21.5%), and (v) 8,308 dispersed duplicates (DSDs, 60.16%) (**Figure 8a, Supplement Table 9**). We then estimated the rates of synonymous and non-synonymous substitutions (*Ks* and *Ka*) for these five gene categories and tested for selection pressures, including positive and purifying selections.

**Figure 8.**
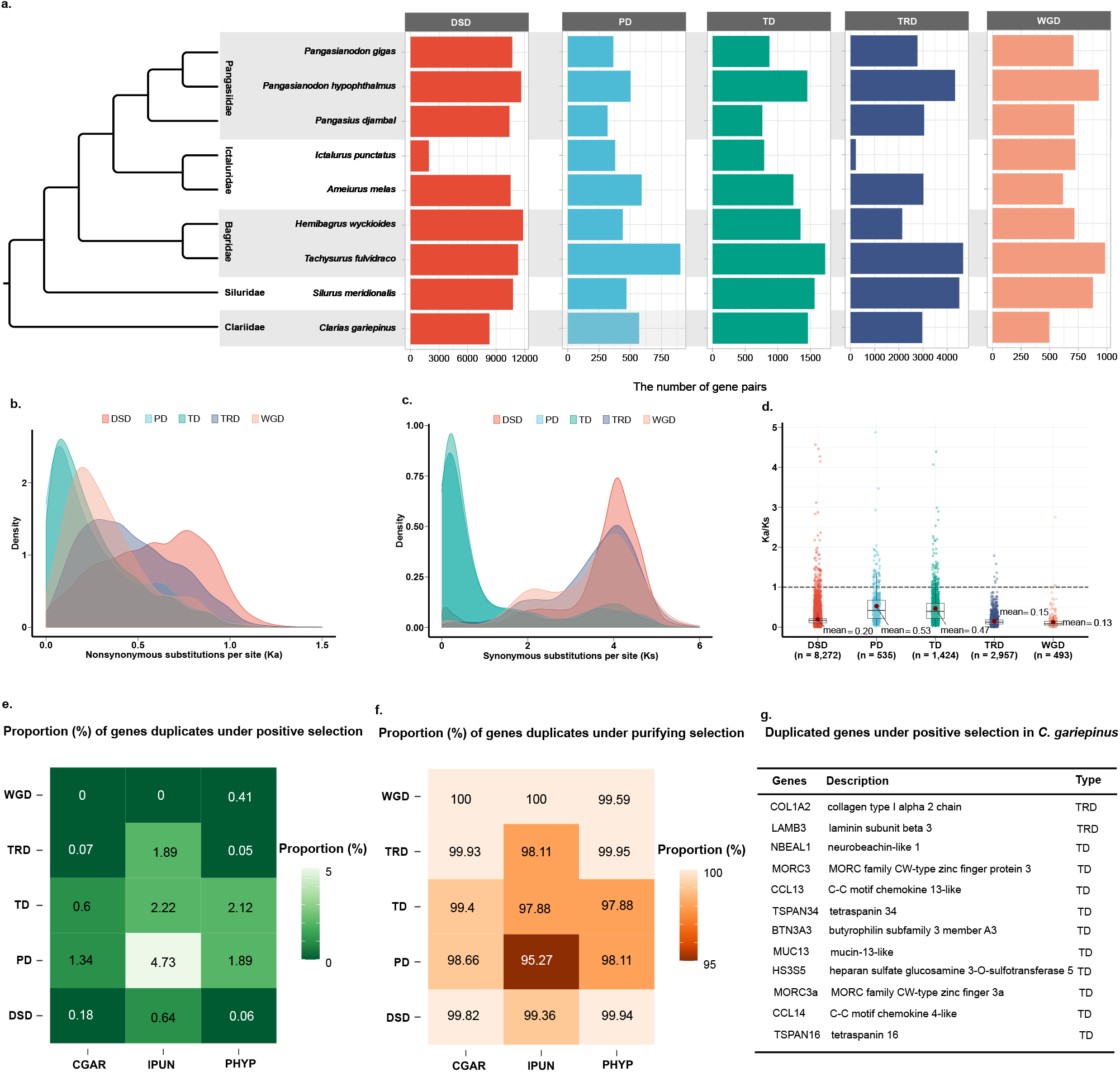
Landscape of gene duplication and positive selection in the African catfish. **a** TThe number of gene pairs derived from various duplication modes in representative catfish genomes. DSD dispersed duplication, PD proximal duplication, TD tandem duplication, TRD transposed duplication, and WGD whole-genome duplication are the different types of duplication. It also shows a schematic representation of the phylogeny of the various catfish species used in the study; **b,c** Evolution of gene pairs duplicated by different modes in African catfish. Ka distributions (**b**) and Ks distributions (**c**); **d** The Ka/Ks ratio distributions of gene pairs derived from different modes of duplication in the African catfish; **e** Percentage of genes under positive selection in three catfish lineages; **f** Percentage of genes under puryfing selection in three catfish lineages. CGAR: The African catfish (*Clarias gariepinus*), IPUN: The channel catfish, (*Ictalurus punctatus*), PHYP: shark catfish (*Pangasius hypophthalmus*); **g** Duplicated genes in *C. gariepinus* that are positively selected in all clariids examined in this study.

The distribution of *Ka*, *Ks*, and *Ka/Ks* among different modes of duplication showed a striking trend, with proximal and tandem duplications having qualitatively higher *Ka/Ks* ratios than other modes. The *Ks* values for PD- and TD-derived gene pairs were relatively low (**Figure 8b-d**). This finding implies that recent TDs and PDs in the African catfish have undergone faster sequence divergence than other paralogs. Whole-genome duplications, on the other hand, are more conserved, with much lower *Ka/Ks* ratios. The three distinct peaks in the *Ks* distribution graph of WGD-derived duplicates reflect the three rounds of teleost-specific WGD, with no evidence of catfish-specific WGD events. All retained WGDs (100%) and nearly all TRDs (99.93%) paralogs were subjected to purifying selection. Positive selection was significantly detected in PDs (1.34%) and TDs (0.6%) duplicate gene pairs. Gene duplications were also analyzed in non-airbreathing catfish species. A similar trend was observed in *Ka/Ks* ratios distribution and the relative proportions of duplicated genes under positive or purifying selection in each paralogs’ category. In particular, purifying selection was observed in 100% and 99.59% of WGD-derived duplicate genes in the channel catfish (*Ictalurus punctatus*, IPUN) and in the shark catfish (*Pangasianodon hypophthalmus*, PHYP), respectively (**Figure 8e-f**). These insights suggest that most duplicated genes were either eliminated or diverged very fast after the most recent whole genome duplication events in catfishes. The generally demonstrated hypothesis of rediploidization substantiates this assumption: the genome tends to return to a stable diploid state by losing one copy of each duplicated gene through non-functionalization and subfunctionalization^109, 110^.

We performed GO enrichment analysis of tandem and proximal duplicates to determine whether the significant selective pressures observed in TDs and PDs drive the evolution of these genes towards biological functions that support the terrestrial adaptation of *Clarias species*. Tandem and proximal-derived duplicates exhibited divergent functional roles although they shared several enriched GO terms involved in immune response, cytoskeletal motor activity, nervous system, and oxygen binding, which are critical for Clarias innate immunity, locomotion and adaptation on land (**Supplement file S2**). In particular, the tandem duplicated mucin-13-like (MUC13) genes are not only under positive selection, but the MUC gene family has also significantly expanded in all five *Clarias* species included in our analysis (**Figure 8g**), suggesting a beneficial or adaptive role for these catfish species.

In summary, these results show that TDs and PDs are substantially involved in the evolutionary mechanisms for adaptation and diversification of airbreathing catfish, as opposed to WGDs and TRDs, which are subjected to strong purifying selection, preventing them from neofunctionalization and subfunctionalization.

## Discussion

Here, we report the first high-quality chromosome-level, haplotype-resolved and near-T2T assembly of the African catfish genome, an economically and ecologically important airbreathing catfish. Leveraging long reads and Hi-C data, we were able to reconstruct the sequences of both haplotypes with total sizes of 954.24 and 972.60 Mbp. Our fully-phased genome assembly exhibited superior quality metrics based on several indicators such as BUSCO, Merqury, phasing accuracy and functional completeness (**Figure 5**, **Table 1**). However, this method only estimates how well each haplotype recovers heterozygous k-mers because only orthogonal reads (e.g. parental) can independently determine the factual phasing accuracy. Although haplotype-specific reads can simulate parental reads, they will still miss a few true heterozygous parental *k*-mers due to sequencing bias or sequencing errors.

We found significant size differences and structural variations between haplotypes in the African catfish genome. Imbalanced assemblies are not uncommon, especially in species with significant size heterogeneity in sex chromosomes. The African catfish (Clarias gariepinus) has a ZZ/ZW sex-determination system in which the sex chromosomes can differ in size. The size heterogeneity of the sex chromosomes may result in variations in haplotype assembly sizes, particularly in sex loci-specific regions. Unfortunately, our data did not allow us to investigate sex-linked variations or definitively identify the sex chromosome in the African catfish genome. However, we conducted validation experiments on previously identified male-specific markers and realized that these markers were present on Hap1 but not on Hap2. This finding suggests that Hap1, the larger haplotype, may represent the male-specific haplotype (including the ZZ male chromosome). To our knowledge, no existing studies so far reported sex chromosome variation or differences in the African catfish to serving as a direct reference for comparison. However, it is worth noting that the observed size difference of around 18 Mb between haplotypes in our study is significantly larger than the expected magnitude of variation observed in a few fish species, such as the goldfish (Carassius auratus), which has a variation of approximately 11.7 Mb^111^. While investigating the structural divergence between haplotypes, we observed that a significant proportion (86.63%) of the 3.5 Mb bases affected by structural variation between Hap1 and Hap2 could be attributed to repeat expansion and contraction (**Figure 9, Supplement Table 10**). Repeat expansion/contraction refers to increased or decreased repeated units within specific genomic regions. This phenomenon likely contributed to the observed size difference between Hap1 and Hap2, potentially leading to underestimating or overestimating the exact gap length between contigs. Collapsed repeats could have further contributed to the size differences, resulting in a shorter assembly for Hap2. In addition, Hap2 was more fragmented than Hap1, as evidenced by a higher number of contigs (212 for Hap2 versus 142 for Hap1) and lower N50 values (19 Mb for Hap2 versus 32 Mb for Hap1). The presence of collapsed repeats and increased fragmentation in Hap2 may have influenced the estimation of overall contiguity and assembly size (**Table 2**).

**Figure 9.**
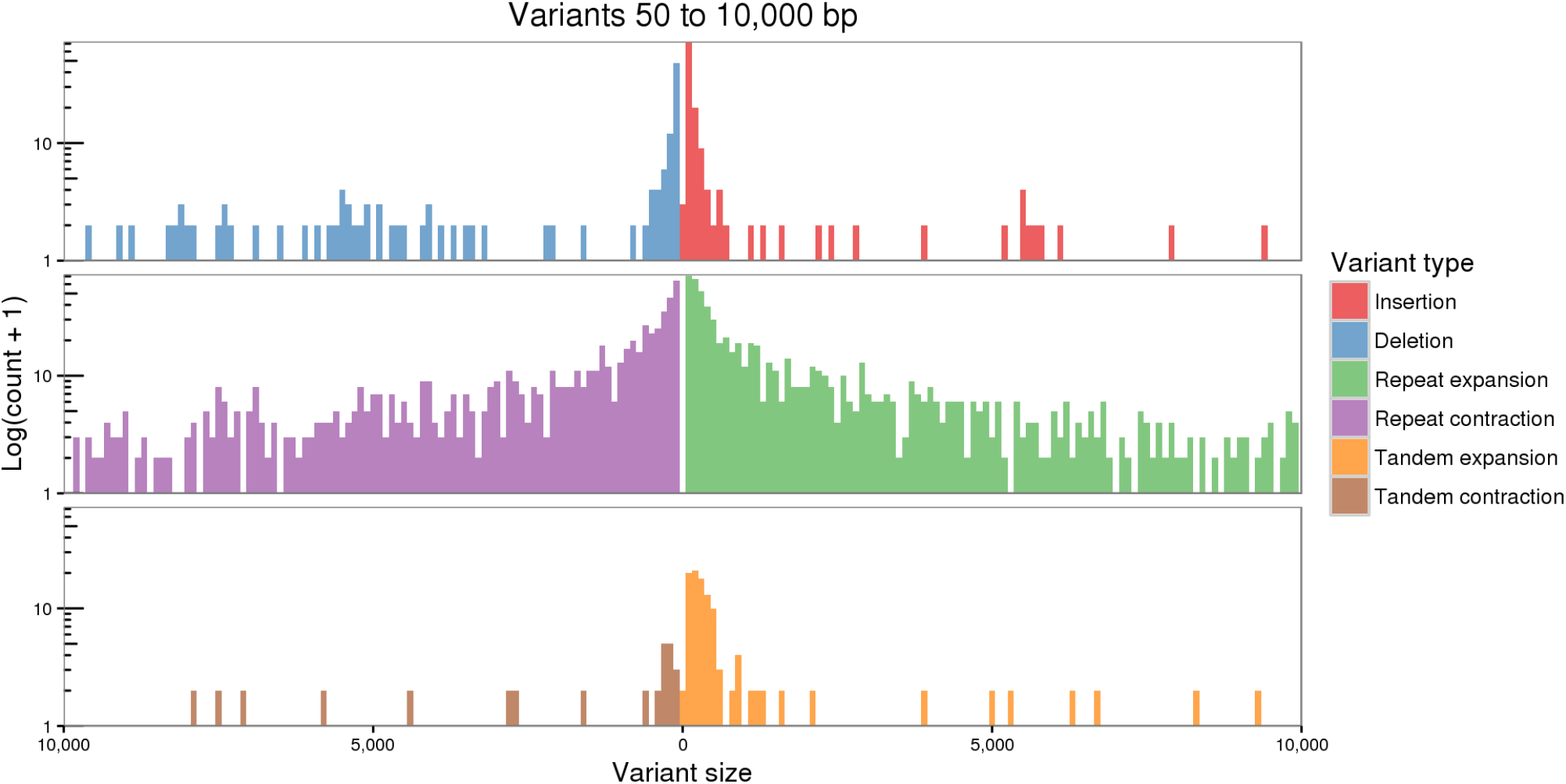
Structural variation between haplotypes. Structural variation (SVs) characterization and quantification between Hap1 and Hap2 using Nucmer and assemblytics. The SVs size range from 50 bp to 10 kbp.

**Table 2.**
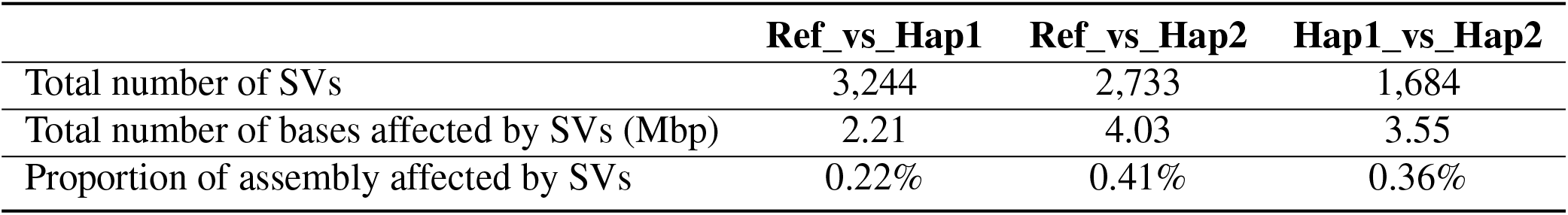
Summary of structural variation quantification between chromosomes of Prim, Hap1 and Hap2 assemblies.

Haplotype-resolved assemblies provide numerous benefits for genomic-based studies of evolution, conservation, and commercial and disease traits. The reported haplotype-resolved genome sequence and annotation provide a powerful tool for enhanced aquaculture and breeding of *C. gariepinus*. It will, for example, aid in sex determination and allow for a better understanding of structural variations, tissue-and haplotype-specific expression. Furthermore, these genomic resources enable more specific investigations of genomic features such as segmental duplications, hybridization, and structural variant hotspots in this and other closely related catfishes^36, 40, 112, 113^. Most (21/28) *C. gariepinus* chromosomes assembly are gapless and resolved from T2T (**Figure 3**). Telomeres are the protective structures that are found at the ends of chromosomes. In teleosts, they consist of a tandemly repeated DNA hexamer (*TTAGGG*)*_n_* and proteins that help to protect the ends of the chromosomes from damage and from being recognized as broken DNA. Our study did not only detect both terminal telomeres in 21 of 28 chromosomes but also several ITS, mainly located at the pericentromeric regions and along the nucleolar organizer regions (NORs). The absence of high-density terminal telomeric signals at both ends of some chromosomes (*n* = 7) is not necessary due to the poor assembly of these regions. The telomeres might be lost or shortened gradually on these chromosomes. The *C. garipinus* genome consists of nine subtelomeric/acrocentric (st/a) chromosomes. It has been established that st/a chromosomes have a very short p-arm and that the length of their telomeres is often shorter than that of other chromosome types^114^. We observed that a few chromosomes without terminal telomeres at both ends exhibit a high copy number of ITS. This suggests that the terminal scaffold is probably misoriented. These ITS may also indicate relics of ancient chromosomal rearrangements in *C. gariepinus*, including centric and tandem chromosome fusion^115, 116^.

Gold standards haplotype-resolved assemblies of commercial fish, such as the one presented here, can aid in designing optimal haplotypes for intra-or interspecies hybridization by avoiding the combination of known incompatibility of alleles. Furthermore, the availability of the two haplotypes of the African catfish is a turning point for modern genomics-assisted breeding strategies for this species. It will ultimately aid in the development of an African catfish breeding program. Our phased assemblies of the African catfish collectively provide the first and most complete view of its genome to date. It paves the way for a variety of applications, including research into the structure and function of telomeres and their role in chromosomal rearrangements and evolution, the loss or fusion of genetic material, and the diversity of karyotypes and sex-chromosome systems in *Claridae*.

Terrestrial adaptation refers to the process by which aquatic species acquire the ability to live or survive on land for an extended period. This process is usually driven by genetic, physiological, and behavioural changes triggered by gene family dynamics, gene duplication events, or positive selection^117, 118^. This evolution can involve many processes and mechanisms, such as changes in body structure, including respiratory and circulatory systems and sensory and nervous systems^102, 119^. Although they acquire certain benefits from the two worlds, bimodal (aerial and aquatic) airbreathing fish face several challenges when adapting to semi-terrestrial habitats. Hypoxic tension, moisture and humidity loss, prolonged exposure to UV radiation, high-temperature fluctuation, locomotion, and exposure to a different spectrum of pathogens, are a few typical challenges that are believed to be the driving forces in the genome remodelling and evolution of aquatic species in these habitats^120–122^. Gene family dynamics (expansion and contraction) is a genome remodelling mechanism that reflects the evolution of organisms’ adaptations to new environments^123^. Our findings show that significantly expanded gene families in Clarias sp. are primarily involved in osmoregulation, anatomical structure development, cytoskeletal motor activity, and stimuli responses. The fluctuating temperature on land will impact osmoregulation and homeostasis via biological processes that regulate ion channels, stress response activation, and osmolyte synthesis^35, 124^. Related gene families such as short-chain dehydrogenases/reductases (SDR), Kv channel interacting proteins (KCNIP), Ferritin (FTH) and hypoxia-inducible factor (EGLN) were significantly associated with these biological processes. We predicted these genes to play a crucial role in the evolution of clariids to semi-terrestrial habitats. For example, FTH plays a role in osmoregulation, particularly in response to changing temperature and salinity^125^. FTH also regulates ion channels and transporters involved in osmoregulation and cells’ adaptation to hypoxic stress^126–128^. In addition, the G-protein-coupled receptor (GPCR) gene family is expanded in the African catfish. Adaptation to terrestrial environments requires fish to maintain proper calcium levels in their bodies as they move between aquatic and terrestrial habitats. Maintaining appropriate calcium levels is crucial for fish on land because calcium is involved in various physiological processes, including muscle contraction, nerve signalling, skeletal development and respiration. These processes may result in structural changes, such as a well-developed fish musculature, that facilitate efficient support and movement on land^3^.

Besides gene family expansion, gene duplication is another process that is believed to trigger the acquisition of evolutionary novelty. It has been reported that gene duplication contributes to the emergence of amphibious traits, which enhance the terrestrial transition of aquatic species^129^. We have characterized recent gene duplication events in selected catfishes, including *C. gariepinus* and other non-airbreathing catfishes. Our findings indicated that TD and PD duplicates exhibited a faster rate of evolution than other modes of duplication, such as WGD, DSD, and TRD. Several TD and PD derived duplicates in *C. gariepinus* were found to be specifically under positive selection in clariids, implying their importance in the differential adaptation of these fish species to new habitats and lifestyles. Gene duplication contributes to gene dosage by increasing the number of genes (gene expansion) that are useful in the adaptation continuum in response to new niches and environments^130, 131^. We found evidence of positive selection in BDH1 (3-hydroxybutyrate dehydrogenase), a member of the SDR protein family. With up to 14 accumulated non-synonymous substitutions, this tandemly duplicated gene showed an accelerated rate of evolution in airbreathing catfishes. Previous studies on *Clarias magur*^132^ and in terrestrial mammals^133^ have found that few members of the SDR gene family, including BDH1, were significantly upregulated in response to low oxygen levels, stressing their potential role in adapting and surviving in hypoxic environments. This is consistent with the hypothesis that airbreathing in fish evolved as a response to aquatic hypoxia^134^.

Overall, these findings suggest that the transition of airbreathing catfish to terrestrial life may rely on a combination of genetic mechanisms such as gene duplication, gene expansion, and positive selection associated with biological processes that shape environmental adaptation. However, it is essential to note that the specific roles of the above-mentioned genes and biological processes in the adaptation of airbreathing catfish remain hypothetical. These predictions lay a solid basis for future studies and further functional validation to fully understand the specific mechanisms that have facilitated the development of additional capabilities for the ecological flexibility of airbreathing catfishes. To fully understand the drivers underlying the adaptation and evolution of this group of fish to terrestrial or semi-terrestrial habitats, extensive research would be needed to establish causal relationships. Undoubtedly this haplotype-resolved assembly, along with the characterization of potential genes and genetic changes/mechanisms involved in environmental adaption, establish the fundamentals for such future studies. These may include studying gene expression patterns in these fish in response to different environmental factors and performing functional validation of these genes’ functions. It could also be insightful to compare the genes and pathways involved in the early evolution and adaption of terrestrial vertebrates to the panel of genes and biological processes hypothesized in this study.

## Conclusions

We have deciphered and annotated the African catfish (*C. gariepinus*) genome, an ecologically and commercially important freshwater airbreathing catfish. This near-T2T chromosome-level assembly and resolved haplotypes advance our understanding of *C. gariepinus* genomic makeup. Comparative genomics analysis with related catfishes provided critical insights into the evolutionary mechanisms underlying airbreathing catfish’s unique terrestrial adaptation, including genes and pathways associated with hypoxia tolerance, locomotion, skeletal muscle development, respiration, osmoregulation, and antioxidant defence. However, to fully uncover the genomic underpinning of these catfishes’ transition from aquatic to terrestrial habitats, further research is needed to validate the specific mechanisms by which these unique genetic changes might have contributed to amphibious traits development in clariids. Furthermore, this work has demonstrated the utility of HiFi data in achieving fully haplotype-resolved genome assemblies. This study provides a valuable resource for future studies on the genomic mechanisms underlying catfishes’ resilience and adaptive mechanisms to adverse ecological conditions. The insights gained could also be leveraged to improve aquaculture practices and enhance the sustainability of catfish farming.

## Data availability

All raw high-throughput sequencing data analysed in this project, including Illumina PE, Hi-C, HiFi, and ONT sequencing reads, are available under NCBI BioProject PRJNA818990. Whole genome assemblies and annotations have been deposited at DDBJ/ENA/GenBank under the accessions GCA_024256425.2 (Primary assembly), GCA_024256435.1 (Haplotype-1) and GCA_024256465.1 (Haplotype-2). The version described in this paper is GCA_024256425.2, GCA_024256435.1, GCA_024256465.1. Further primary data and information on research design are provided at Zenodo (`10.5281/zenodo. 7760650`).

## Code availability

Customs scripts and pipelines used in the data analysis and to create figures are available at https://github.com/ bbalog87/catfish-genome

## Acknowledgements

We thank Dr. Alexander Rebl for his constructive comments and input towards interpreting the results.

## Funding

This work was funded by the European Maritime and Fisheries Fund (EMFF). EMFF grant: MV-II.1-LM-014

## Author contributions

T.G., R.M.B. and J.A.N. conceptualized the project. T.G. acquired funding. R.M.B. and J.A.N. collected and prepared the tissue samples for sequencing. J.A.N., Y.A.B.Z., T.G. and R.M.B. designed the methodology. J.A.N. and Y.A.B.Z. performed whole bioinformatics analyses and developed the figures. J.A.N. wrote the original draft manuscript. T.G., R.M.B., Y.A.B.Z. and J.A.N. contributed to reviewing and editing the manuscript. All authors read and approved the submitted version.

## Competing interests

The authors declare no competing interests.

## Notes

https://zenodo.org/record/7760650

### Competing Interest Statement

The authors have declared no competing interest.

### Summary of Updates

We have perfomed additional valiadtion of the data and extended the discussion section.

